# Brain DNA Methylation Patterns in *CLDN5* Associated With Cognitive Decline

**DOI:** 10.1101/857953

**Authors:** Anke Hüls, Chloe Robins, Karen N. Conneely, Rachel Edgar, Philip L. De Jager, David A. Bennett, Aliza P. Wingo, Michael P. Epstein, Thomas S. Wingo

## Abstract

**Objective:** Cognitive decline is a hallmark of dementia; however, the brain epigenetic signature of cognitive decline is unclear. We investigated the associations between brain tissue-based DNA methylation and cognitive trajectory.

**Methods:** We performed a brain epigenome-wide association study of cognitive trajectory in 636 participants from the Religious Order Study and the Rush Memory and Aging Project (ROS/MAP) using DNA methylation profiles of the dorsal lateral prefrontal cortex (dPFC). To maximize our power to detect epigenetic associations, we used the recently developed Gene Association with Multiple Traits (GAMuT) test to analyze the five measured cognitive domains simultaneously.

**Results:** We found an epigenome-wide association for differential methylation of sites in the Claudin-5 (*CLDN5*) locus and cognitive trajectory (p-value x 9.96 × 10^-7^), which was robust to adjustment for cell type proportions (p-value = 8.52 x 10^-7^). This association was primarily driven by association with declines in episodic (p-value = 4.65 x 10^-6^) and working memory (p-value = 2.54 x 10^-7^). This association between methylation in *CLDN5* and cognitive decline was independent of beta-amyloid and neurofibrillary tangle pathology and present in participants with low levels of neuropathology. In addition, only 13-31% of the association between methylation and cognitive decline was mediated through levels of neuropathology, whereas the major part of the association was independent of it.

**Interpretation:** We identified methylation in *CLDN5* as new epigenetic factor associated with cognitive trajectory. Higher levels of methylation in *CLDN5* were associated with faster cognitive decline implicating the blood brain barrier in maintenance of cognitive trajectory.

## Introduction

Cognitive decline is a common concern among older adults; however, the trajectory of cognitive performance with age has a wide range from stable to rapid decline. Cognitive trajectory is an important predictor of health outcomes and mortality, independent of other commonly assessed risk factors^1^. Dementia is a common consequence of a decline in cognition, and Alzheimer Disease (AD) is its leading cause^2^. AD is characterized by the neuropathological accumulation of neuritic plaques and neurofibrillary tangles, which is accompanied by neuronal loss^3^; however, most older individuals have several co-occurring neuropathologies. Collectively, neuropathologies explain about 40% of the variance in cognitive trajectory, leaving most unexplained^4,5^. Thus, cognitive trajectory may be considered a summation of the different neuropathological and biological processes independent of pathologies at work in the aging human brain^5–7^.

Despite the importance of understanding cognitive trajectory, existing epigenetic work on DNA methylation levels measured in brain tissue has primarily focused on AD-specific pathologies ^8–11^ and clinical diagnosis of AD ^12–14^. In contrast, epigenetic studies that focused on examining cognitive decline were limited due to measuring DNA methylation changes in blood^15^, which showed only moderate correlations (~0.4) with brain methylation^15^. Thus, there is need to understand the epigenetic changes that are associated with cognitive trajectory to identify potential mechanisms that may act through or independent of known neuropathologies.

In this study, we investigated the associations between brain tissue-based DNA methylation and cognitive trajectory in 636 participants from the Religious Order Study and Rush Memory and Aging Project (ROS/MAP) cohorts. Cognitive trajectory was assessed in five cognitive domains (episodic memory, perceptual speed, perceptual orientation, semantic memory, and working memory), which were analyzed simultaneously by using an innovative kernel procedure that allows to investigate associations between multiple predictors (e.g. methylation sites in a gene) with multiple outcomes (e.g. multiple cognitive domains) ^16,17^. Findings were validated in independent post-mortem frontal cortex samples^10^, and their biological plausibility were evaluated using gene expression and genotype data.

## Methods

### Study design and study population

The discovery dataset included deceased subjects from two large, prospectively followed cohorts recruited by investigators at Rush Alzheimer’s Disease Center in Chicago, IL: The Religious Orders Study (ROS) and the Rush Memory and Aging Project (MAP)^11,18^. Both ROS and MAP collect detailed annual cognitive and clinical evaluations, and brain autopsy. Participants provided informed consent, an Anatomic Gift Act for organ donation, and a repository consent to allow their data to be repurposed. Both studies were approved by an Institutional Review Board of Rush University Medical Center. To be included in the present study, participants must have at least two follow-up evaluations, and available methylation data derived from dorsolateral prefrontal cortex. As in previous publications, the ROS and MAP data were analyzed jointly since much of the phenotypic data collected are identical at the item level in both studies and collected by the same investigative team^11,19^.

The replication dataset included samples from the MRC London Neurodegenerative Disease Brain Bank (GSE59685)^10^.

### DNA methylation

In the discovery dataset, DNA methylation was measured from the dorsolateral prefrontal cortex (dPFC; Broadman area 46) as previously described in 737 ROS/MAP participant samples ^11^, of which 665 had complete phenotype and covariate information. DNA was extracted from cortically dissected sections of dPFC and DNA methylation was measured using the Illumina HumanMethylation450 Beadchip array. Initial data processing, including color channel normalization and background removal, was performed using the Illumina GenomeStudio software. The raw IDAT files were obtained from Synapse (www.synapse.org; Synapse ID: syn7357283) and the following probes were removed: 1) probes with a detection p-value > 0.01 in any sample, 2) probes annotated to the X and Y chromosomes by Illumina, 3) probes that cross-hybridize with other probes due to sequence similarity (identified by ^20^), 3) non-CpG site probes, and 4) probes that overlap with common SNPs (identified by ^21^). After this filtering, the remaining CpG sites were normalized using the BMIQ algorithm in Watermelon R package^22^, and the ComBat function from the sva R package was used to adjust for batch effects^23^. CpG sites with a distance of more than 20 KB to the closest gene were excluded from analysis. After quality control 338,036 discrete CpG dinucleotides corresponding to 26,558 genes in 636 subjects were used for analysis.

In the replication dataset, DNA methylation was derived from prefrontal cortex obtained from individuals archived in the Medical Research Counsil (MRC) London Neurodegenerative Disease Brain Bank. This previously published dataset provided samples with DNA methylation data measured on the Illumina HumanMethylation450 Beadchip array. Data was obtained from the Gene Expression Omnibus (GEO; GSE59685)^10^. Similar to above probes annotated to the X and Y chromosomes, cross-hybridizing probes, non-CpG site probes and probes that overlap with common SNP were removed. The data downloaded from GEO was already normalized using the dasen algorithm in Watermelon R package^22^ and then the ComBat function from the sva R package was used to adjust for array ID batch effects ^23^. Neuronal and non-neuronal brain cell-type proportions were estimated and normalized between PFC samples^24^. CpG sites with a distance of more than 20 KB to the closest gene were excluded from analysis. After quality control 358,515 discrete CpG dinucleotides corresponding to 27,585 genes in 66 subjects were used for analysis.

### Genotype data

Genotyping data was generated using two microarrays, Affymetrix GeneChip 6.0 (Affymetrix, Inc, Santa Clara, CA, USA) and Illumina HumanOmniExpress (Illumina, Inc, San Diego, CA, USA) as described previously^25^. Genotyping was imputed to the 1000 Genome Project Phase 3 using the Michigan Imputation Server ^26^, and the following filtering criteria were applied minor allele frequency (MAF) > 5%, Hardy-Weinberg p-value >10^-5^ and genotype imputation R^2^ > 0.3.

### Gene expression

RNA extracted from ROS/MAP post-mortem dPFC was sequenced on the Illumina HiSeq with 101-bp paired-end reads using the strand-specific dUTP method with poly-A selection with a coverage of 50 million reads. BAM files were converted to FASTQ format using Picard, followed by alignment of reads to GRCh38 reference genome using STAR^27^. Gene level counts were computed using STAR^27^. Genes with < 1 count per million in at least 50% of the samples and with missing length and percent GC content were removed. Additionally, two outlier samples were removed. After quality control, counts were normalized using variance stabilization transformation, which performed log2 transformation of the counts, normalizes for library size, and transforms the counts to approximately homoscedastic^28^. Then the candidate mRNAs were extracted for association analysis with rate of cognitive decline adjusting for sex, age at death, RIN, PMI, RNA-sequencing batch, and cell type composition. Proportions of neurons, astrocytes, oligodendrocytes, and microglia were estimated from RNA-sequencing data using CIBERSORT^29^ and cell-type specific signatures^30^. We used the proportions of cell type to adjust for tissue heterogeneity. Using the findings of the epigenome-wide association study of cognitive trajectory, we selected the transcripts corresponding to the associated genes for further analyses after the aforementioned quality control and variance stabilization transformation.

### Cognitive trajectory

Cognitive trajectory was assessed in five different cognitive domains: episodic memory, perceptual speed, perceptual orientation, semantic memory, and working memory. Participants in both studies underwent structured, annual clinical evaluations that included detailed cognitive and neurologic examinations, as previously reported ^31,32^. Scores from those tests were converted to z-scores using the mean and standard deviation of the cohorts at baseline. Cognitive scores were modeled longitudinally with a mixed effects model, adjusting for age, sex and education, providing person-specific random slopes of decline. The random slope of each subject captures the individual rate of cognitive decline after adjusting for age, sex and education.

### Neuropathologic Outcomes

We used the CERAD score and Braak staging as neuropathological outcomes in our analyses. The CERAD score is a semiquantitative measure of neuritic plaque density as recommended by the Consortium to Establish a Registry for Alzheimer’s Disease (CERAD). A CERAD neuropathological diagnosis of AD requires moderate (probable AD) or frequent neuritic plaques (definite AD) in one or more neocortical regions. The Braak stage is a standardized measure of neurofibrillary tangle distribution and burden determined at autopsy with modified Bielschowsky silver stained sections^33^. Braak stages I and II indicate neurofibrillary tangle confined mainly to the entorhinal region of the brain, Braak stages III and IV indicate involvement of limbic regions such as the hippocampus and Braak stages V and VI indicate moderate to severe neocortical involvement.

### Statistical analysis

In our main analysis, we estimated epigenetic associations across five neurocognitive domains in a gene-based analysis, in which each CpG site was assigned to the closest gene using the Bioconductor package hiAnnotator^34^ and the ensembl gene predictions (ensGene, version of Apr-06-2014). All CpG sites with a distance of no more than 20 KB to the closest gene, were included in the analyses. In addition, we conducted a sensitivity analysis in which we only included CpG sites with a distance of no more than 10 KB to the closest gene. In a traditional association study of cognitive trajectory, each cognitive test may be either tested individually ^15^ or used to estimate a composite measure aggregated across several cognitive tests^35^. However, these approaches are underpowered in the presence of pleiotropy since they fail to exploit correlation among domains^17^. Thus, analysis of a single composite measure can lose power if the causal CpG sites are only associated with a subset of the features that make the composite measure^17^. Hence, it can be more powerful to directly account for the trait correlations using kernel methods^36^. Kernel methods quantify the genetic similarity among pairs of subjects and test whether this genetic similarity is associated with trait similarity. Thus, they harness potential pleiotropy that exists between traits to improve power to detect associations. To analyze epigenetic associations across five neurocognitive domains simultaneously, we applied a variation of the GAMuT test ^16^ that was adapted to DNA methylation data. GAMuT is motivated by the idea that individuals with similar epigenetic patterns should also have similar cognitive traits across the different cognitive domains. Consequently, GAMuT constructs two different similarity matrices; one similarity matrix including cognitive decline in the five cognitive domains and the other similarity matrix for the epigenetic variation (beta values of CpG sites) assigned to a gene. Phenotypic and epigenetic similarity were modelled using linear kernels. P-values for GAMuT were derived using Davies’ exact method, which is a computationally efficient method to provide accurate p-values in the extreme tails of tests that follow mixtures of chi-square variables ^37,38^. To test which cognitive domains and CpG sites were likely main drivers in our multivariate analysis, we conducted an association analysis for each domain separately. The gene-based analyses for the single domains were performed with GAMuT and linear regression analyses were used in the CpG-based analyses.

All association models were adjusted for age at death, education, sex, ancestry, smoking status and post-mortem interval (PMI). Principal components (PCs) based on CpG sites chosen for their potential to proxy nearby SNPs (within 10 BP) were used to correct for population stratification (first three PCs, Figure S1) and cell type heterogeneity ^21^. Samples whose first PC (PC1) deviated more than 3 standard deviations from the mean PC1, were excluded from analyses, reducing the final sample size from 665 to 636. In a sensitivity analysis, associations were additionally adjusted for cell type proportions^24^. All analyses were performed using R (version 3.4.3) using built-in functions unless otherwise specified.

We applied a Bonferroni threshold to correct for multiple testing. In the gene-based GAMuT analysis, the significance threshold was adjusted for the number of tested genes (threshold: 0.05/26,558 = 1.88 x 10^-6^) and in the CpG-site-based linear regression analyses for the number of tested CpG sites (threshold: 0.05/338,036=1.48 x 10^-7^).

Methylation signals associated with cognitive decline were validated using CERAD and Braak stage as outcomes. In addition, we analyzed whether the association between methylation and cognitive decline was modified (interaction analysis) or mediated (causal mediation analysis) by neuropathology (CERAD, Braak stage). Causal mediation analysis from a counterfactual perspective was performed by using the R package “mediation” ^39^, an approach that relies on the quasi-Bayesian Monte Carlo method based on normal approximation ^40^. Using the counterfactual framework allows for definition of direct and indirect effects and a total effect as the sum of direct and indirect effects. The indirect effect refers to the effect through the mediator under study. The direct effect refers to the remaining effect that is not through the mediator ^41^. The proportion of the indirect effect in the total effect was used to assess the extent to which the association between methylation and cognitive decline was mediated through neuropathology as an intermediate pathway ^42^.

The replication dataset did only have Braak stage as neurocognitive outcome, which is strongly associated with cognitive decline ^33^. In the replication dataset, associations between CpG sites within a gene and the Braak stage were tested with GAMuT after correction for age at death, sex and cell type composition. Due to the small sample size of the replication dataset, we conducted a permutation test with 10,000 replications in addition to the Davies’ approximation to verify the accuracy of p-values.

The biological plausibility of our findings was examined by investigating the association 1) of DNA methylation with gene expression as well as of gene expression with cognitive decline and 2) of genotypes with cognitive decline to investigate if our associations were due to a hidden genotype effect. All of these associations were tested using GAMuT and the genotype associations were followed by a linear regression analysis on the single SNP level. Fine-mapping of our epigenome-wide associations was done with coMET ^43^, which is a visualization tool of EWAS results with functional genomic annotations and estimation of co-methylation patterns.

## Results

### Description of study participants

There were 636 ROS/MAP participants included in this study with an average age at death of 86 years and with 63% being female (Table 1). Most of the participants were white (98%), had a high level of education and 70% had never smoked. On average, cognitive performance declined with age for every single domain (Table 1) and correlations of cognitive decline between different domains were moderate, ranging from 0.54 to 0.78 (Table S1). Cognitive decline was associated with more signs of neuropathology (CERAD and Braak stage, Table S2).

**Table 1.** Study characteristics of the discovery (ROS/MAP) and replication datasets.

|  | Discovery Dataset | Replication Dataset |
| --- | --- | --- |
| N | 636 | 66 |
| Age at death, mean ± sd | 86.22 ± 4.72 | 87.48 ± 6.57 |
| Female, n (%) | 401 (63.1%) | 44 (66.67%) |
| Ancestry |  |  |
| European, n (%) | 621 (97.6%) | *n.a.* |
| African American, n (%) | 11 (1.7%) | *n.a.* |
| Native American, n (%) | 1 (0.2%) | *n.a.* |
| Asian, n (%) | 3 (0.5%) | *n.a.* |
| Years of education, mean ± sd | 16.63 ± 3.54 | *n.a.* |
| Never smoker, n (%) | 444 (69.8%) | *n.a.* |
| Ex-smoker, n (%) | 176 (27.7%) | *n.a.* |
| Smoker, n (%) | 16 (2.5%) | *n.a.* |
| Post mortem interval (PMI), mean ± sd | 7.43 ± 5.79 | *n.a.* |
| Decline in episodic memory, mean ± sd | -0.03 ± 0.11 | *n.a.* |
| Decline in perceptual speed, mean ± sd | -0.02 ± 0.08 | *n.a.* |
| Decline in perceptual orientation, mean ± sd | -0.01 ± 0.04 | *n.a.* |
| Decline in semantic memory, mean ± sd | -0.03 ± 0.13 | *n.a.* |
| Decline in working memory, mean ± sd | -0.01 ± 0.05 | *n.a.* |
| CERAD, mean ± sd | 2.27 ± 1.15 | *n.a.* |
| Braak stage, mean ± sd | 3.44 ± 1.26 | 4.70 ± 1.62 |

*n.a.*: information not available in replication dataset.

There were 66 MRC Brain Bank participants with an average age at death of 87 years, with 67% being female and an average Braak stage of 5 (Table 1).

### Methylation patterns of CLDN5 associated with cognitive decline

In the ROS/MAP participants (discovery dataset), we found that methylated CpG sites in the Claudin-5 (*CLDN5*) locus were associated with cognitive trajectory (p-value = 9.96 x 10^-7^; Figure 1, Table 2, Table S3, and Figure S1). This association was robust to adjustment for cell type proportions (p-value = 8.52 x 10^-7^, Table S4, Figures S2 and S3), to the selection of a smaller window of CpG sites around each gene (p-value = 9.96 x 10^-7^, within 10kb, Table S5), and to the restriction to participants with European ancestry (621/636 participants, p-value = 9.73 x 10^-7^, Table S6). The trajectories of episodic and working memory were the main drivers for the observed association with both being associated with CpG sites assigned to *CLDN5* in the analyses of the single domains (Table 2, Figure S4 and Tables S7-S11). Genes showing suggestive association with cognitive trajectory (p-values < 5 x 10^-5^) included *AC084018.1, CTB-186G2.1, ATG16L2, KCNN4, RP11-779O18.1, TTC22, DCUN1D2-AS, PNMA1 and RP11-101C11.1*. The strongest associations with these genes were found with episodic memory, followed by working memory. Interestingly, most top methylation signals in Table 2 (*CLDN5* and 7/9 suggestive genes) were also at least nominally associated with CERAD (Table 3, Figures S5-S6 and Table S12) and Braak stage (Table 3, Figure S7 and Table S13).

**Figure 1.**
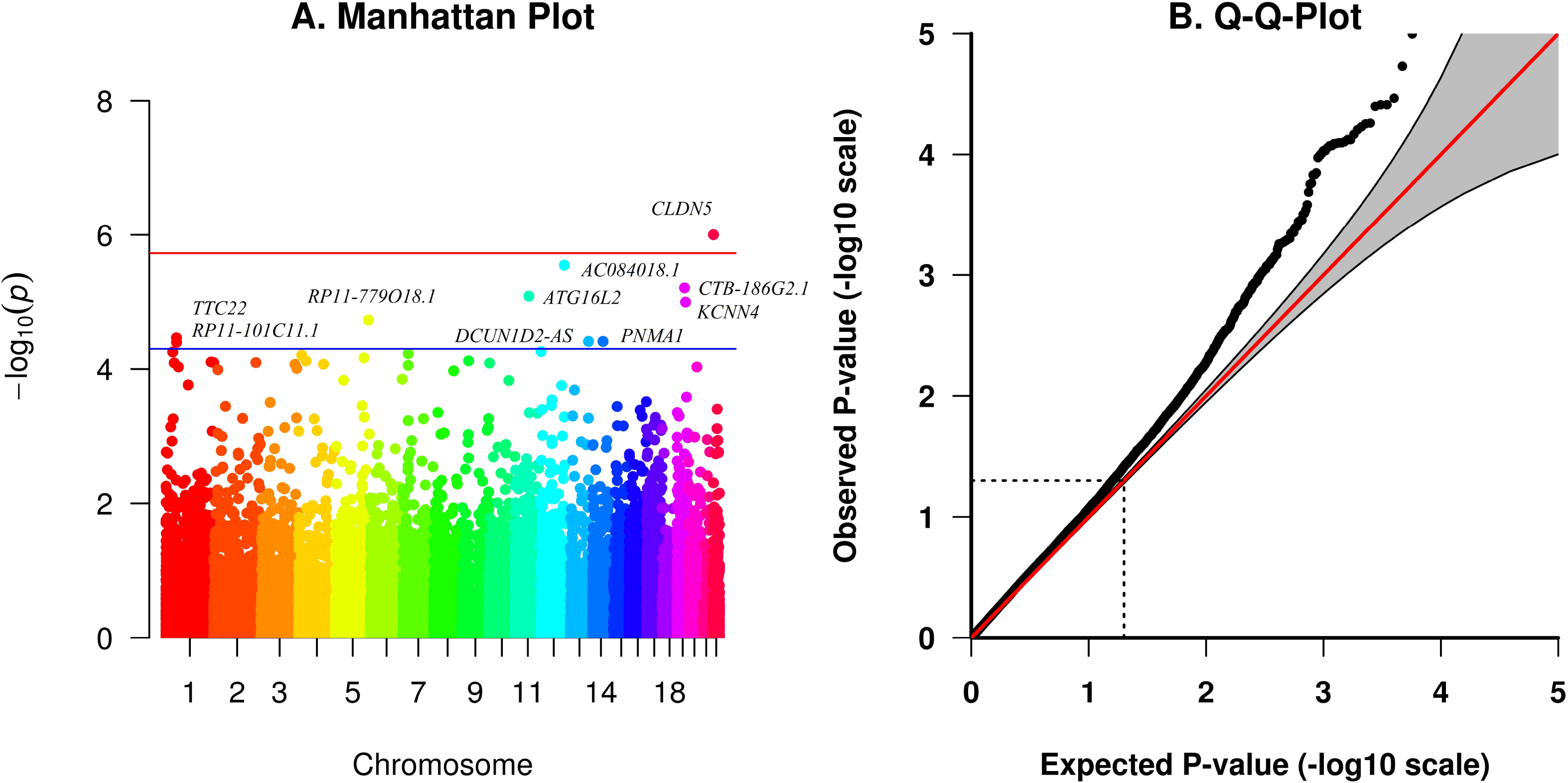
DNA methylation and cognitive decline. Association between DNA methylation and cognitive decline in ROS/MAP (discovery dataset) tested with GAMuT. Adjusted for age at death, education, sex, ancestry, smoking status, post-mortem interval (PMI) and the first three principal components.

**Table 2.** Top signals (p-values < 5 x 10^-5^) for the association between DNA methylation and cognitive decline across all domains as well as in the single domains in the discovery dataset (ROS/MAP).

| gene | chr | pos (lower) | pos (upper) | mean<br>dist gene | max<br>dist gene | #CpG sites | p-value<br>(across all<br>domains) | p-value<br>(EM) | p-value<br>(PS) | p-value<br>(PO) | p-value<br>(SM) | p-value<br>(WM) |
| --- | --- | --- | --- | --- | --- | --- | --- | --- | --- | --- | --- | --- |
| CLDN5 | chr22 | 19510877 | 19515608 | 1030 | 2062 | 18 | **9.96E-07** | <u>4.65E-06</u> | 7.16E-05 | 0.0021 | <u>2.35E-05</u> | **2.54E-07** |
| AC084018.1 | chr12 | 122235169 | 122241475 | 501 | 1414 | 15 | <u>2.84E-06</u> | 5.96E-05 | 0.0001 | 0.0060 | <u>4.13E-06</u> | 0.0002 |
| CTB-186G2.1 | chr19 | 39087135 | 39090701 | 692 | 1633 | 4 | <u>6.19E-06</u> | <u>7.33E-06</u> | <u>3.50E-05</u> | 0.0015 | 7.45E-05 | 7.69E-05 |
| ATG16L2 | chr11 | 72521478 | 72546168 | 1326 | 12094 | 25 | <u>8.22E-06</u> | <u>1.32E-05</u> | 0.0002 | 0.0086 | 8.31E-05 | 0.0004 |
| KCNN4 | chr19 | 44270892 | 44286076 | 1682 | 6854 | 13 | <u>1.01E-05</u> | <u>2.01E-05</u> | 0.0002 | 0.0127 | 0.0001 | <u>4.12E-05</u> |
| RP11-779O18.1 | chr5 | 172168177 | 172189374 | 7685 | 17051 | 12 | <u>1.86E-05</u> | <u>1.26E-05</u> | 0.0002 | 0.0262 | 0.0009 | <u>3.18E-06</u> |
| TTC22 | chr1 | 55240609 | 55268659 | 1189 | 4775 | 23 | <u>3.42E-05</u> | 0.000192 | 0.0002 | 0.0182 | 0.0002 | 6.16E-05 |
| DCUN1D2-AS | chr13 | 114123258 | 114129580 | 2530 | 3583 | 10 | <u>3.87E-05</u> | <u>5.91E-06</u> | 0.0004 | 0.0176 | 0.0014 | <u>4.48E-05</u> |
| PNMA1 | chr14 | 74177136 | 74181427 | 777 | 1357 | 9 | <u>3.88E-05</u> | <u>3.14E-05</u> | 0.0079 | 0.0030 | 0.0002 | 0.0006 |
| RP11-101C11.1 | chr1 | 55682652 | 55709508 | 5270 | 9658 | 2 | <u>3.99E-05</u> | <u>2.36E-05</u> | 0.0004 | 0.0565 | 0.0006 | 0.0003 |

EM: decline in episodic memory; PS: decline in perceptual speed; PO: decline in perceptual orientation; SM: decline in semantic memory; WM: decline in working memory
Adjusted for age at death, education, sex, ancestry, smoking status, post-mortem interval (PMI) and the first three principal components (PCs).
Bonferroni threshold: $0.05/26,558=1.88 \times 10^{-6}$ (p-values below Bonferroni threshold in **bold**)
Suggestive: p-values $< 5 \times 10^{-5}$ (underlined)

**Table 3.** Validation of ROS/MAP (discovery dataset) findings in replication dataset

| gene | chr | pos (lower) | pos (upper) | Discovery Dataset |  |  |  | Replication Dataset |  |  |
| --- | --- | --- | --- | --- | --- | --- | --- | --- | --- | --- |
|  |  |  |  | #CpG sites | p-value cognitive decline | p-value CERAD | p-value Braak | #CpG sites | p-value Braak | p-value Braak (permutation Test<sup>1</sup>) |
| CLDN5 | chr22 | 19510877 | 19515608 | 18 | 9.96E-07 | **0.0023** | **0.0066** | 18 | **0.0088** | **0.0076** |
| AC084018.1 | chr12 | 122235169 | 122241475 | 15 | 2.84E-06 | **0.0002** | **5.31E-06** | 16 | 0.1794 | 0.1791 |
| CTB-186G2.1 | chr19 | 39087135 | 39090701 | 4 | 6.19E-06 | **0.0005** | **8.36E-05** | 4 | **0.0125** | **0.0121** |
| ATG16L2 | chr11 | 72521478 | 72546168 | 25 | 8.22E-06 | **0.0003** | **5.67E-06** | 25 | 0.1535 | 0.1536 |
| KCNN4 | chr19 | 44270892 | 44286076 | 13 | 1.01E-05 | **0.0028** | **0.0001** | 15 | **0.0222** | **0.0178** |
| RP11-779O18.1 | chr5 | 172168177 | 172189374 | 12 | 1.86E-05 | **0.0168** | **0.0008** | 12 | 0.0890 | 0.0872 |
| TTC22 | chr1 | 55240609 | 55268659 | 23 | 3.42E-05 | **0.0010** | **0.0002** | 24 | 0.2027 | 0.2096 |
| DCUN1D2-AS | chr13 | 114123258 | 114129580 | 10 | 3.87E-05 | **0.0363** | **0.0344** | 11 | 0.6988 | 0.7057 |
| PNMA1 | chr14 | 74177136 | 74181427 | 9 | 3.88E-05 | 0.0709 | 0.1913 | 10 | 0.1022 | 0.0999 |
| RP11-101C11.1 | chr1 | 55682652 | 55709508 | 2 | 3.99E-05 | 0.1052 | 0.0831 | 2 | 0.1582 | 0.1654 |

Discovery dataset: ROS/MAP samples. All analyses were adjusted for age at death, education, sex, ancestry, smoking status, post-mortem interval (PMI) and the first three principal components (PCs). Associations with CERAD were additionally adjusted for cell type proportions and the fourth PC.
Replication dataset: MRC Brain Bank Samples (Lunnon et al., 2014) All analyses were adjusted for age at death, sex and cell type composition.
Successful replication (p-value < 0.05 in validation cohort) in **bold**.
<sup>1</sup>Due to the small sample size (N=66) we conducted a permutation test with 10,000 replications in addition to the Davies' approximation to verify the p-values in the validation cohort.

To generalize our findings from the ROS/MAP participants, we performed the same epigenetic analysis in MRC Brain Bank participants with DNA methylation data available (replication dataset). We tested for differential methylation using Braak stage. Of the 8 methylation signals, which were at least nominally associated with Braak stage in the discovery dataset, *CLDN5, CTB-186G2.1* and *KCNN4* could be replicated in the replication dataset (Table 3, Table S14, Figure S8).

### Higher levels of methylation in CLDN5 locus associated with cognitive decline

To identify which CpG sites are the main drivers of the observed associations and to understand the direction of association, we conducted a linear regression analysis for the cognitive trajectory of each cognitive domain. Interestingly, except for *PNMA1*, higher levels of methylation within our top genes were associated with an increased cognitive decline in every single cognitive domain (Table S15). Within the CpG sites assigned to *CLDN5*, cg16773741 and cg05460329 were the main drivers of the association with cg16773741 being associated with episodic memory (p-value = 1.48 x 10^-8^), semantic memory (8.81 x 10^-8^) and working memory (8.66 x 10^-9^) (Table 4). The direction of association with these two CpG sites was consistent with the association observed for CERAD and Braak stage, which could further be replicated in the replication dataset (Table S15, N=66).

**Table 4.** Identification of lead signals within *CLDN5* and direction of associations.

| gene | chr | pos | CpG site | beta<br>(EM) | p-value<br>(EM) | beta<br>(PS) | p-value<br>(PS) | beta<br>(PO) | p-value<br>(PO) | beta<br>(SM) | p-value<br>(SM) | beta<br>(WM) | p-value<br>(WM) |
| --- | --- | --- | --- | --- | --- | --- | --- | --- | --- | --- | --- | --- | --- |
| CLDN5 | chr22 | 19512903 | cg05460329 | -0.93 | 1.43E-06 | -0.62 | 1.60E-05 | -0.25 | 0.0003 | -1.10 | 1.89E-06 | -0.45 | 2.41E-06 |
| CLDN5 | chr22 | 19512942 | cg05498726 | -0.55 | 0.0001 | -0.36 | 0.0008 | -0.13 | 0.0162 | -0.61 | 0.0005 | -0.26 | 0.0002 |
| CLDN5 | chr22 | 19513006 | cg11450827 | -0.60 | 0.0007 | -0.35 | 0.0064 | -0.16 | 0.0111 | -0.65 | 0.0019 | -0.30 | 0.0005 |
| CLDN5 | chr22 | 19513008 | cg17583256 | -0.57 | 0.0005 | -0.30 | 0.0129 | -0.14 | 0.0180 | -0.51 | 0.0086 | -0.27 | 0.0008 |
| CLDN5 | chr22 | 19513017 | cg16773741 | -0.76 | **1.48E-08** | -0.39 | 8.38E-05 | -0.18 | 0.0003 | -0.86 | **8.81E-08** | -0.38 | **8.66E-09** |
| CLDN5 | chr22 | 19513078 | cg14553765 | -0.44 | 0.0014 | -0.32 | 0.0017 | -0.13 | 0.0087 | -0.60 | 0.0003 | -0.29 | 1.92E-05 |
| CLDN5 | chr22 | 19513176 | cg00189989 | -0.61 | 0.0004 | -0.42 | 0.0009 | -0.14 | 0.0224 | -0.78 | 0.0001 | -0.36 | 1.86E-05 |

Only CpG sites with a p-value < 0.001 for at least one cognitive domain are shown. Bonferroni threshold: 0.05/338,036=1.48e-07 (p-values below Bonferroni threshold in bold). EM: decline in episodic memory; PS: decline in perceptual speed; PO: decline in perceptual orientation; SM: decline in semantic memory; WM: decline in working memory. Adjusted for age at death, education, sex, ancestry, smoking status, post-mortem interval (PMI) and the first four principal components.

### Association with CLDN5 even present without signs of neuropathology

To investigate if the association between methylation in CLDN5 and cognitive decline is also present in participants without clear signs of neuropathology, we conducted an analysis of the interaction between the most significantly associated CpG site (cg16773741) and CERAD or Braak stage on cognitive decline. The association between methylation in cg16773741 did not significantly differ between participants with no to little signs of neuropathology versus participants with moderate to severe signs of neuropathology (measured by CERAD and Braak stage; Figure 3). Consequently, the association between methylation in cg16773741 and decline in episodic, semantic, and working memory was even significant in participants with no or little signs of neuropathology.

**Figure 2.**
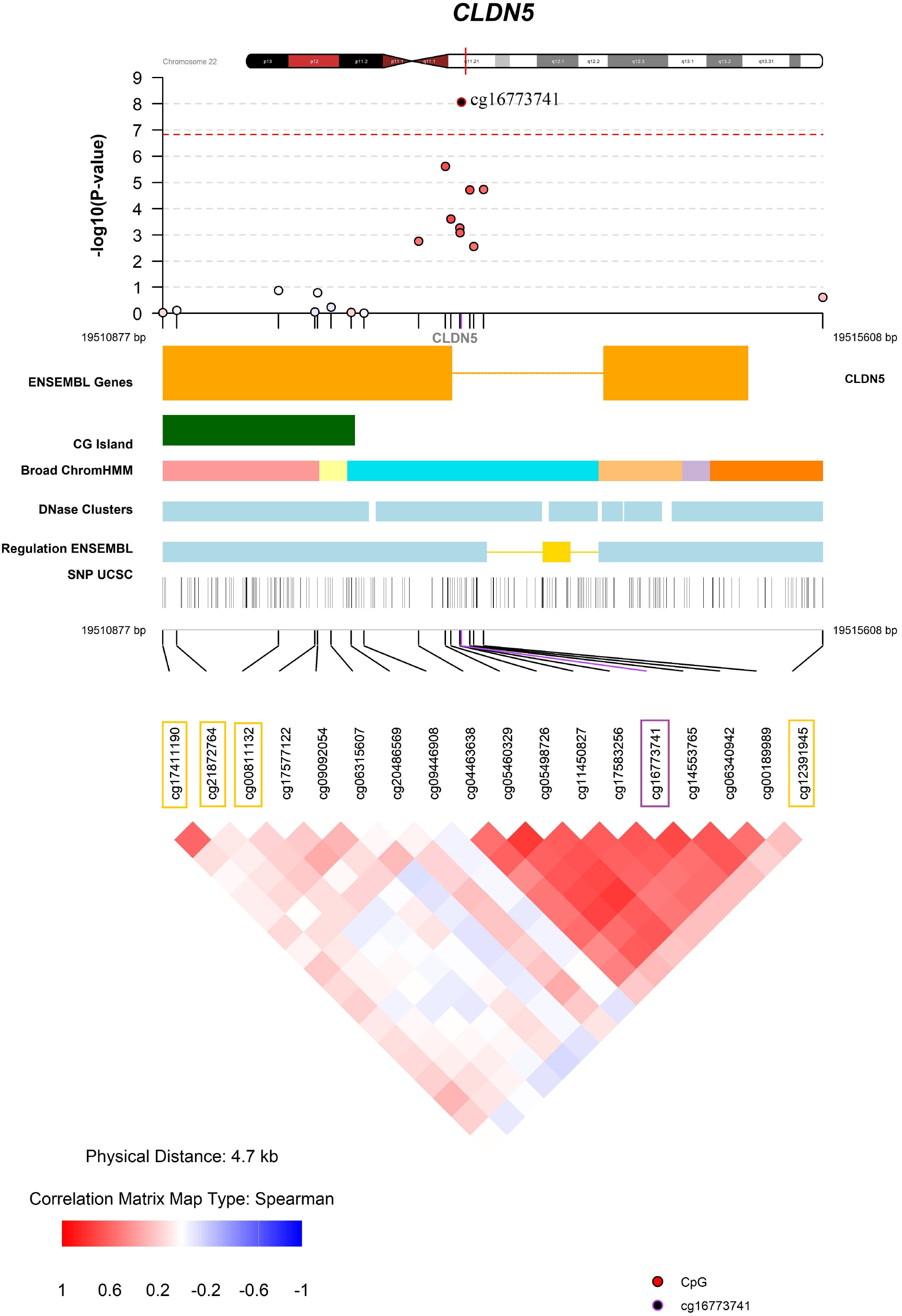
Fine mapping of the association between DNA methylation and decline in working memory. Results from linear regression analyses on the association between CpG sites and cognitive decline in ROS/MAP (discovery dataset) adjusted for age at death, education, sex, ancestry, smoking status, post-mortem interval (PMI) and the first four principal components. The most significant CpG site (cg16773741) is marked in purple and CpG sites associated with genotypes in the same window are marked in yellow (compare Table S17).

**Figure 3.**
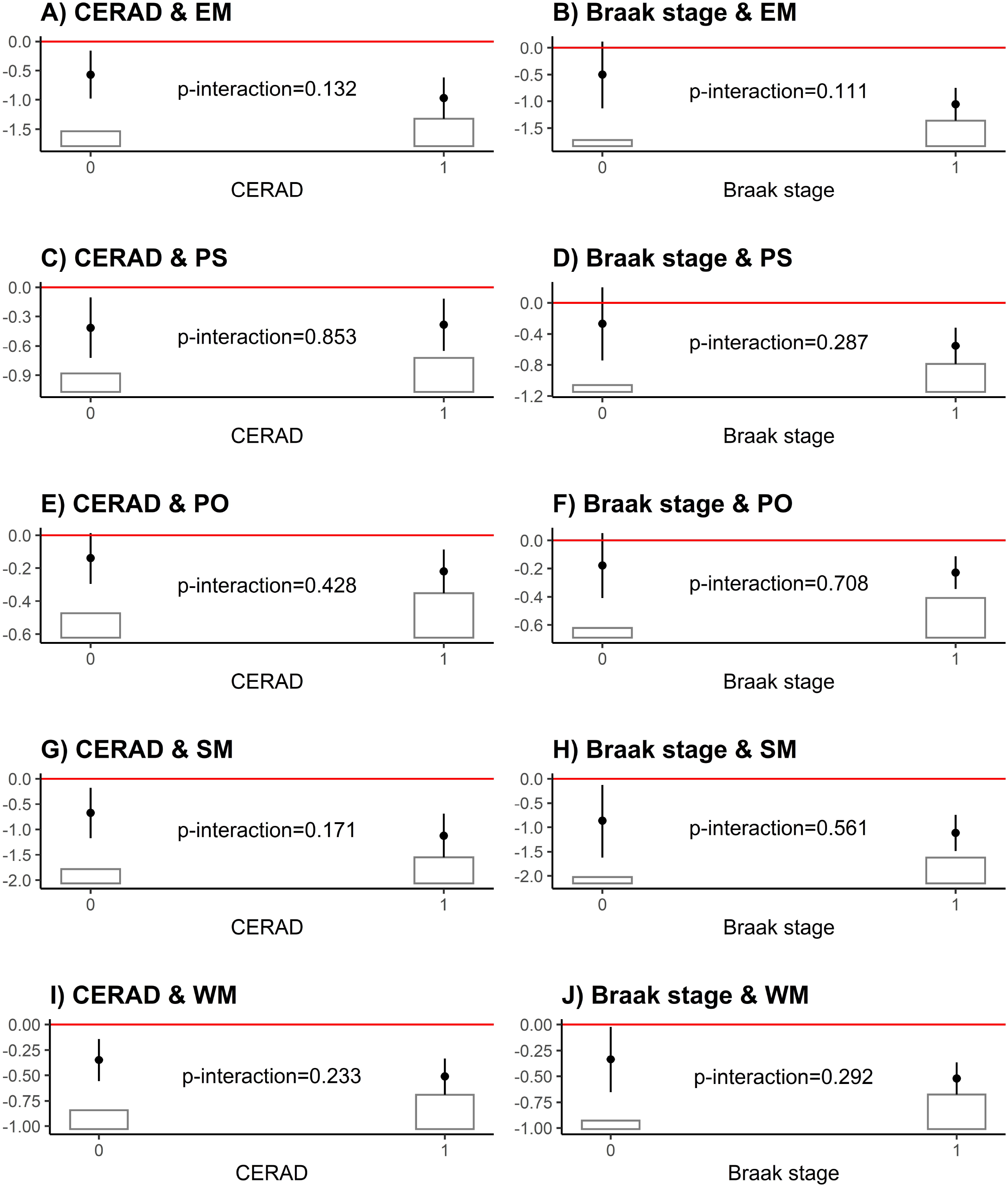
Interaction analysis. Associations between DNA methylation of the top CpG site of *CLDN5* (cg16773741) and cognitive trajectory are shown in participants with no to little (category 0) vs. moderate to severe (category 1) signs of neuropathology. No to little signs of neuropathology are defined as a CERAD measure of 3 (possible) or 4 (no AD) or a Braak stage of 0 to II. Moderate to severe signs of neuropathology are defined as a CERAD measure of 1 (definite) or 2 (probable) or a Braak stage of III to VI. P-values are given for the test of deviations of the association between methylation in cognitive trajectory between the two strata. The bars present the distribution of the neuropathological variables. EM: decline in episodic memory; PS: decline in perceptual speed; PO: decline in perceptual orientation; SM: decline in semantic memory; WM: decline in working memory. Adjusted for age at death, education, sex, ancestry, smoking status, post-mortem interval (PMI) and the first three principal components.

### Partial mediation through neuropathology

The association between DNA methylation in the *CLDN5* locus (cg16773741) and cognitive trajectory was only partially mediated through an increased neuropathology. It ranged between 17% (95% confidence interval (CI): 18-40%) to 31% (95% CI: 18-61%) for CERAD and 13% (95% CI: 7-21%) to 27% (95% CI: 15-41%) for Braak stage depending on the cognitive domain (Figure 4). Therefore, the major part of the association with *CLDN5* was a direct association between methylation and cognitive decline, which was independent of beta-amyloid and neurofibrillary neuropathology.

**Figure 4.**
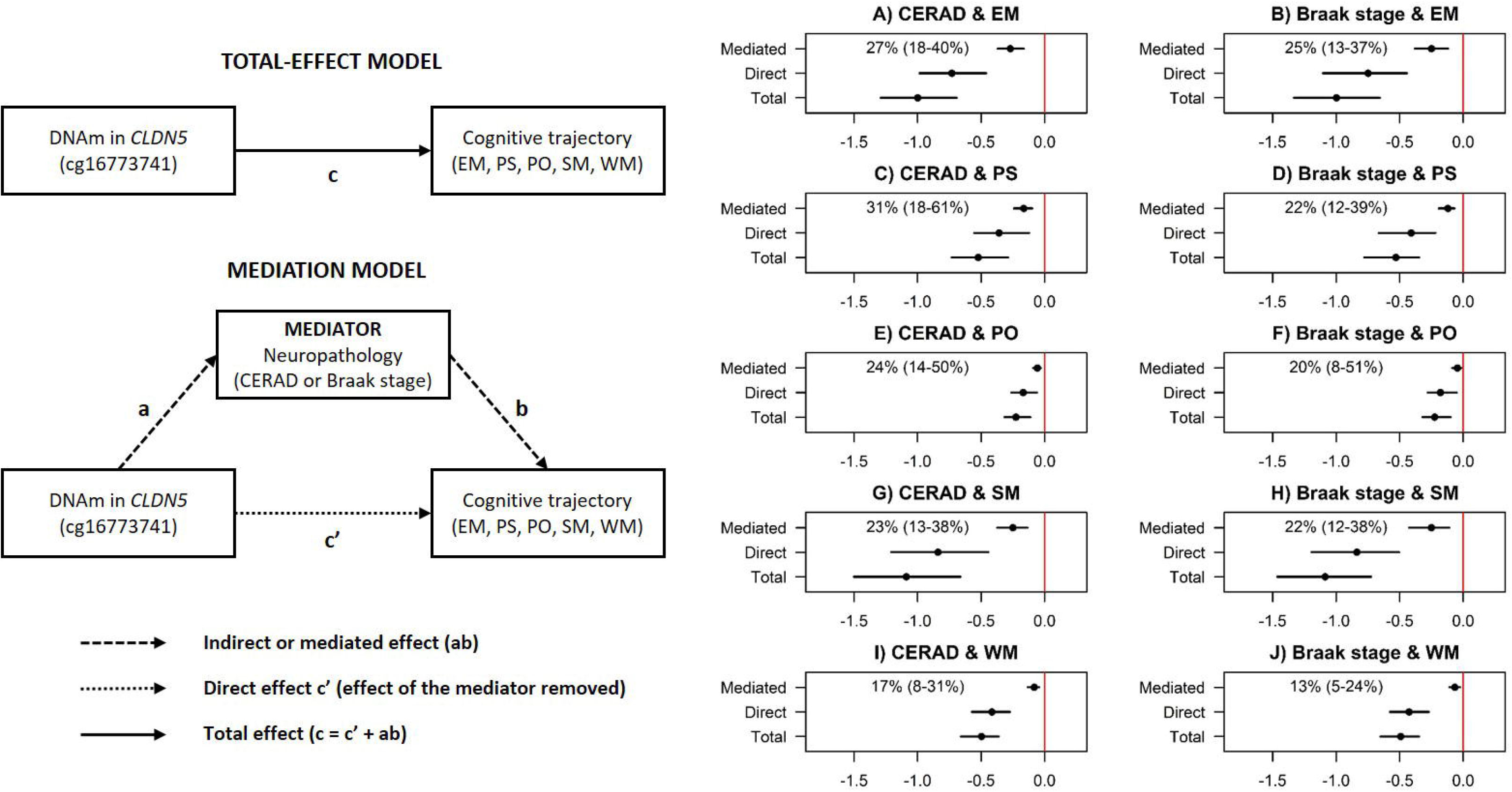
Causal mediation analysis. Beta-estimates and 95%-confidence intervals of the estimated average causal mediation effects, the average direct effects as well as the total effects. Proportion (with 95%-confidence interval) of the association between DNA methylation (DNAm) of the top CpG site of *CLDN5* (cg16773741) and cognitive trajectory, which is mediated through neuropathology (CERAD & Braak stage) is given in percent. EM: decline in episodic memory; PS: decline in perceptual speed; PO: decline in perceptual orientation; SM: decline in semantic memory; WM: decline in working memory. Adjusted for age at death, education, sex, ancestry, smoking status, post-mortem interval (PMI) and the first three principal components (PCs).

### Methylation signals were independent of genotypes

Genotypes located within the windows of *CLDN5* were associated with DNA methylation (p-value < 10^-6^), but not with cognitive trajectory (p-value = 0.4415, Table S16 A). In line, associations of DNA methylation in the *CLDN5* window with cognitive trajectory were robust to adjustment for genotypes from the same window (Table S16 B).

This observation was confirmed by the subsequent analyses on the single CpG / SNP level, which showed that the CpG sites associated with genotypes were not the same as being associated with cognitive trajectory (Table S17, Figure 2 and Figures S9 to S12). This indicates that our methylation signals were not caused by hidden genotype effects.

### No clear association with gene expression levels

In our sample, we find no association between DNA methylation in the *CLDN5* window and *CLDN5* expression (p-value = 0.1978; Table S17). *KCNN4* was the only top gene (Table 2) for which methylation levels were associated with expression levels (p-value = 0.0004; Table S18). Furthermore, cognitive trajectory was not associated with expression of any gene in Table 2 (Table S18).

## Discussion

In this study, we found an epigenome-wide association between brain-tissue-based DNA methylation in the *CLDN5* locus and cognitive trajectory in more than 600 participants from the ROS/MAP cohort. This association was significant across different domains and particularly associated with trajectories in episodic and working memory. We also found that higher levels of methylation in *CLDN5* were associated with neuropathology in our discovery and replication datasets consistent with the direction of association found with cognitive decline. Most interestingly, the association between methylation in CLDN5 and cognitive decline was independent of beta-amyloid and neurofibrillary neuropathology and even present in participants with low levels of those pathologies. In addition, only 13-31% of the association between methylation and cognitive decline was mediated through levels of neuropathology, whereas the major part of the association was independent of it. Finally, we found no evidence that hidden effects of genotypes in the *CLDN5* locus confounded our methylation results.

*CLDN5* is an integral membrane protein and an important component of tight junction protein complexes that comprise the blood-brain barrier. The blood-brain barrier is located at endothelial cells lining the brain microvasculature and is maintained by the neurovascular unit, a functional relationship between astrocytes, neurons, and endothelial cells ^44^. Dysfunction of the blood-brain barrier has been implicated in neurodegenerative disorders, such as AD ^44–47^. Thus, our finding that altered regulation of *CLDN5* is associated with cognitive decline suggests a role of blood-brain barrier dysfunction in cognitive decline. We note that an estimated two-thirds of AD dementia (clinically defined) and that an estimated 40% of cognitive decline are attributable to known age-related neuropathologies ^4,48^. Thus, our findings may account for some of the unaccounted-for variation in cognitive decline and AD dementia.

This is the first epigenome-wide study using cognitive trajectory in older individuals. A previous study on the same cohort showed an association of 71 CpG sites with neuritic plaque burden, of which 11 were validated in an independent cohort^11^. Here, we showed that all of these 11 signals were at least nominally associated with cognitive trajectory and the strongest associations were again found for cognitive trajectory of episodic and working memory (Table S19). In addition, we identified methylation in *CLDN5* as new epigenetic factor associated with cognitive trajectory, a gene that has not been linked to AD in a population-based cohort.

By contrast, associations of blood-based methylation levels with global cognitive function (cg21450381) and phonemic verbal fluency (cg12507869) ^15^ could not be validated in our study (Table S20). The likely reasons are the different source of methylation data, and differences in the phenotype (i.e., cognitive trajectory over time in our study versus cognitive testing at a single time point^15^).

Strengths of this study include the ROS/MAP cohort as a discovery dataset which is notable for its longitudinal nature with very high follow-up rates, prospective collection of data, a community-based cohort design, and detailed neuropathological examination following high autopsy rates. The validity of our signals was also determined in an independent replication dataset, and the availability of genomic data allowed us to determine methylation changes were not the result of hidden genotype effect. This study is also strengthened by the GAMuT ^16^ statistical method that harnesses correlations among cognitive domains and among CpG sites to improve statistical power compared to standard univariate techniques.

The study is potentially limited by its cross-sectional nature. Although brain tissue is the ideal target tissue to measure DNA methylation related to cognitive trajectory, it inhibited a simultaneous (or even later) assessment of cognitive function. Consequently, we cannot exclude the potential risk of reverse causality in our associations. Another potential limitation is the use of bulk tissue analysis which might obscure signals from different cell populations. This problem was mitigated in our analysis by adjusting for cell-type composition. However, the bulk tissue analysis may have obscured an association between *CLDN5* methylation and RNA expression. Future studies should investigate the role of *CLDN5* in specific cell types from brain and investigate whether there is a causal relationship between *CLDN5* dysregulation and cognitive decline in animal models of AD.

In conclusion, we have presented evidence for brain-based DNA methylation in association with cognitive trajectory. We identified methylation in *CLDN5* as a new epigenetic factor associated with cognitive trajectory, which was validated in an independent dataset and independent of beta-amyloid and neurofibrillary neuropathology. Higher levels of methylation in *CLDN5* were associated with cognitive decline implicating the blood brain barrier in maintenance of cognitive trajectory with aging.

## Acknowledgments

The authors are grateful to the participants of the Rush Memory and Aging Project and Religious Orders Study and the Medical Research Counsel Brain Bank.

## Author Contributions

AH planned and conducted the analyses supported by MPE, KC, TSW and APW. AH, TSW, APW and MPE were the major contributors in writing the manuscript. KC and CR were responsible for the preprocessing and quality control of the DNA methylation data from the discovery cohort and RE for the replication cohort. APW conducted the preprocessing and quality control of the RNAseq data and TSW was supervised the quality control of the genotype data. PLDJ generated the methylation, transcriptomic, and genetic data. DAB conceived, designed, and leads the ROS/MAP study and the methylation sub-study.

## Web resources

Data used in this study: https://www.synapse.org/#!Synapse:syn2580853

Rush Alzheimer’s Disease Center Research Resource Sharing Hub: www.radc.rush.edu.

Epstein Software: https://github.com/epstein-software.

MRC London Neurodegenerative Diseases Brain Bank: https://www.kcl.ac.uk/ioppn/depts/bcn/our-research/neurodegeneration/brain-bank.

Ensembl gene predictions (ensGene, version of Apr-06-2014): http://hgdownload.soe.ucsc.edu/goldenPath/hg19/database/.

