## Supplementary Figures for "Brain DNA Methylation Patterns in *CLDN5* Associated With Cognitive Decline"

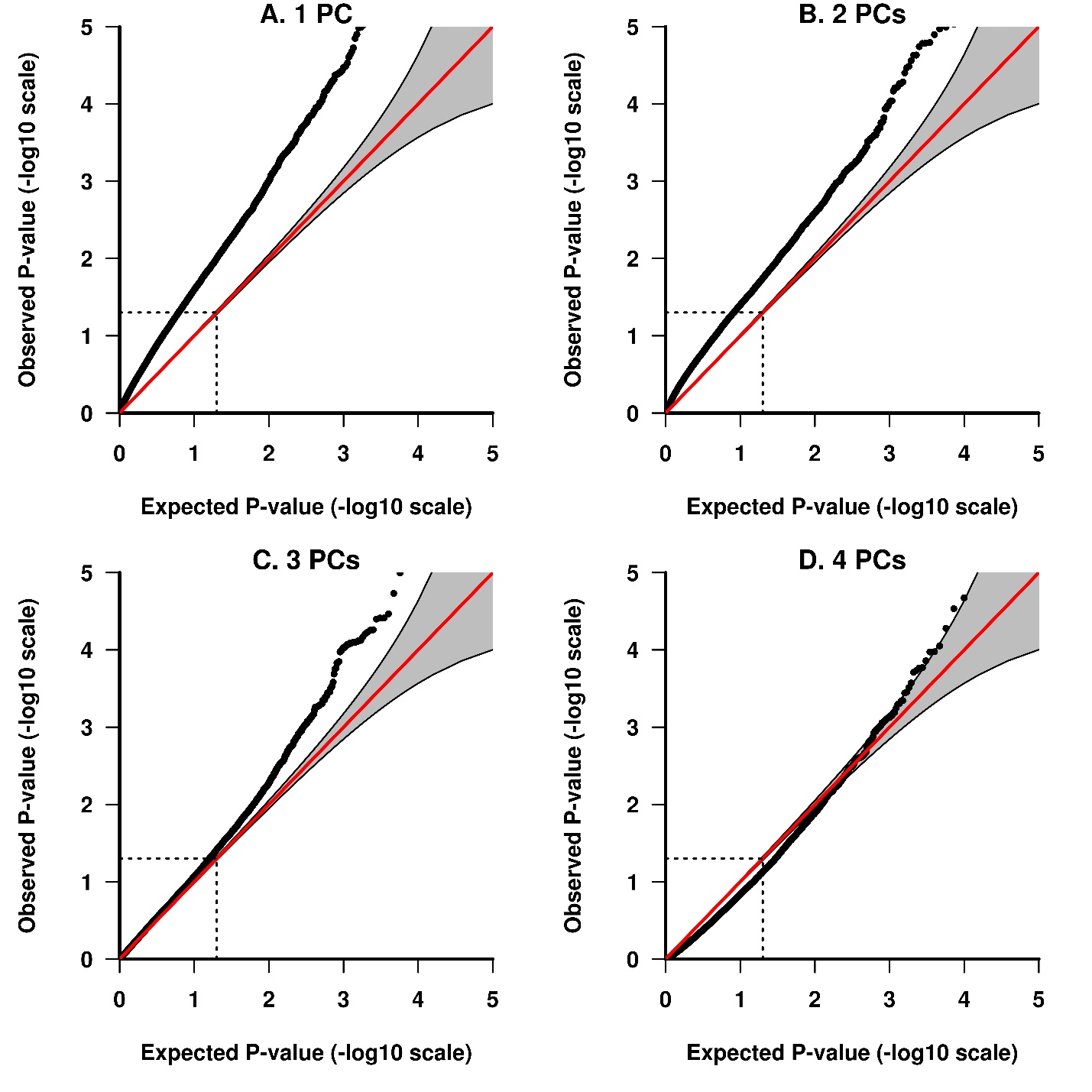

**Figure S1.** Impact of number of principal components (PCs) in the adjustment set on the type I error. Association between DNA methylation and cognitive decline in ROS/MAP (discovery dataset) tested with GAMuT. Adjusted for age at death, education, sex, ancestry, smoking status, post-mortem interval (PMI) and the first 1 to 4 principal components.

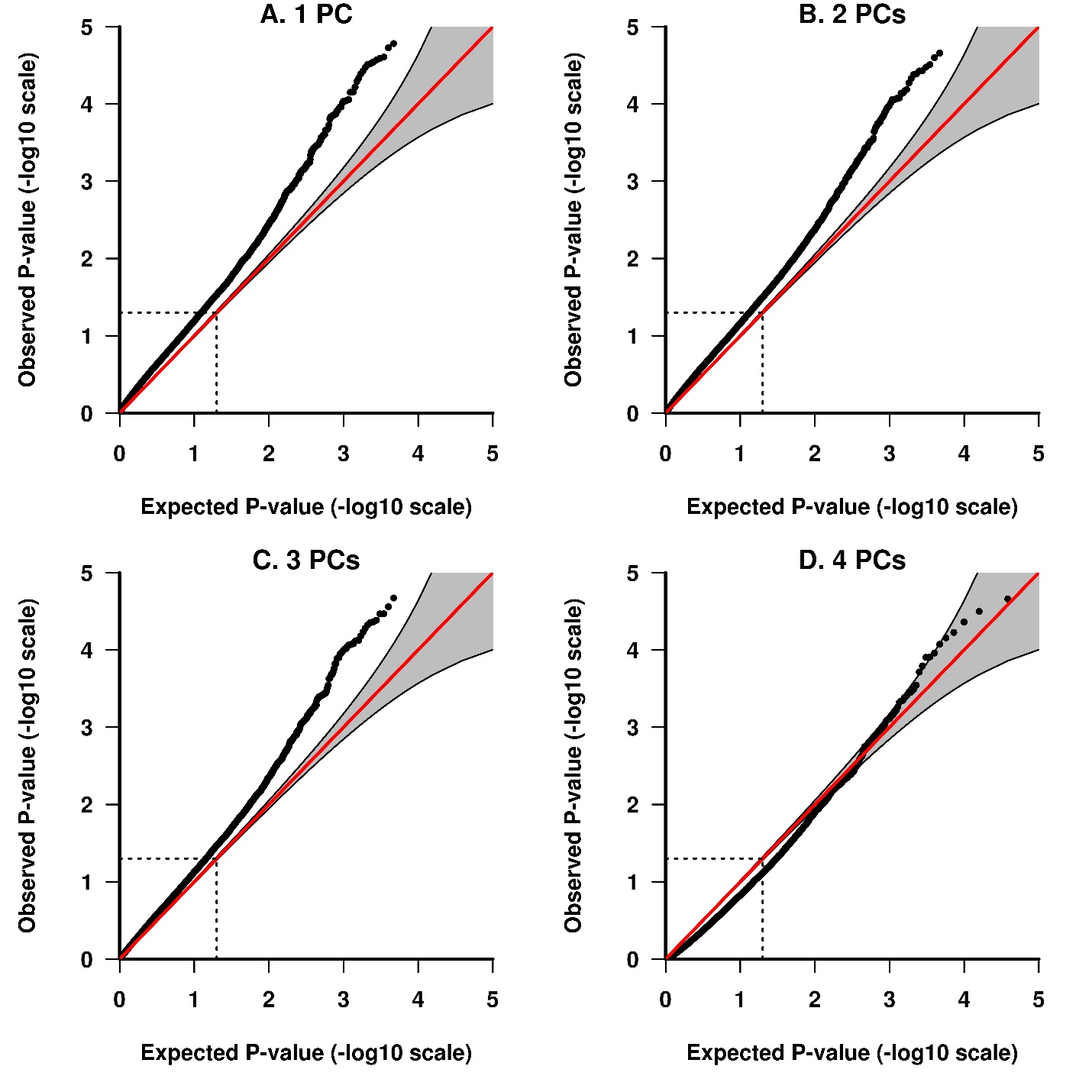

**Figure S2.** Impact of number of principal components (PCs) in the adjustment set on the type I error. Association between DNA methylation and cognitive decline in ROS/MAP (discovery dataset) tested with GAMuT. Adjusted for age at death, education, sex, ancestry, smoking status, post-mortem interval (PMI), cell type proportions and the first 1 to 4 principal components.

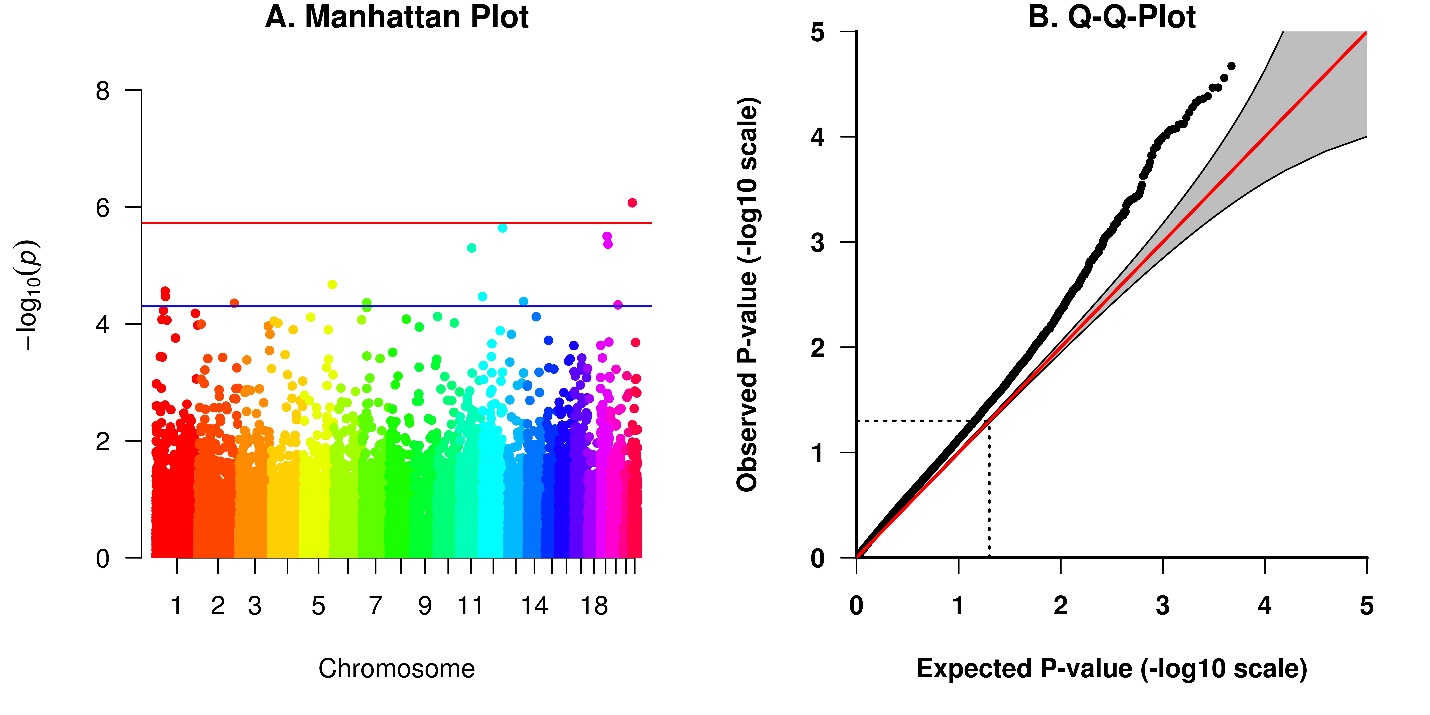

**Figure S3.** Association between DNA methylation and cognitive decline in ROS/MAP (discovery dataset) tested with GAMuT. Adjusted for age at death, education, sex, ancestry, smoking status, post-mortem interval (PMI), cell type proportions and the first three principal components.

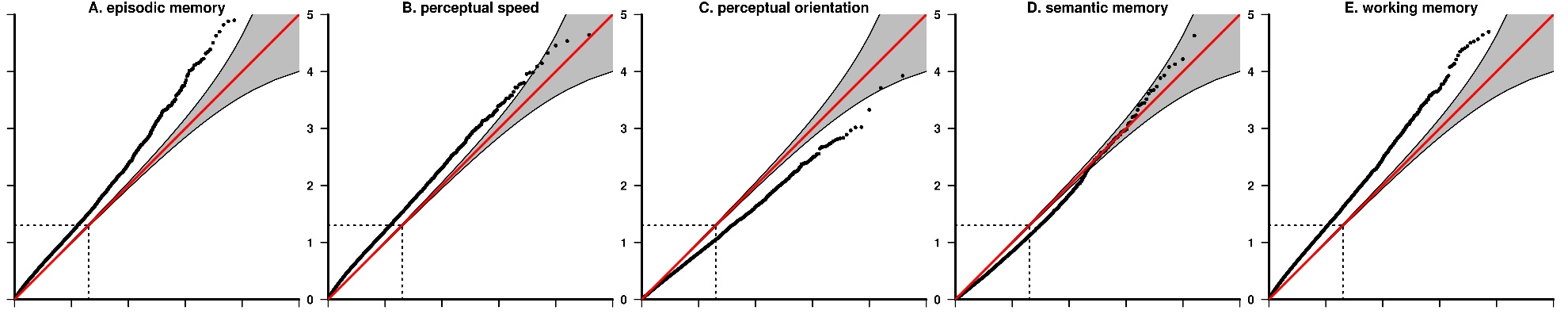

**Figure S4.** Association between DNA methylation and cognitive decline in the single domains in ROS/MAP (discovery dataset) tested with GAMuT. Adjusted for age at death, education, sex, ancestry, smoking status, post-mortem interval (PMI) and the first 3 principal components.

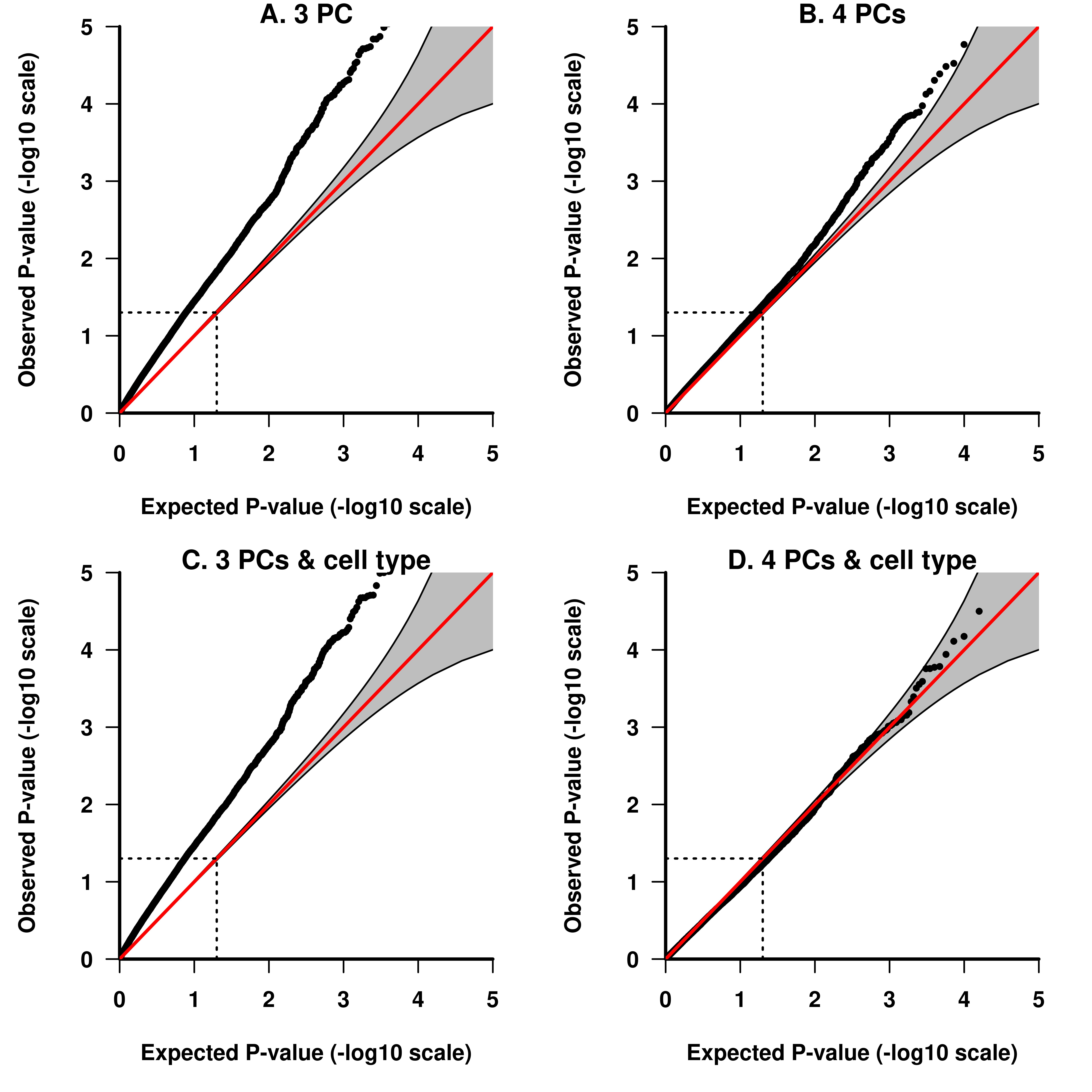

**Figure S5.** Impact of the adjustment set on the type I error. Association between DNA methylation and CERAD in ROS/MAP (discovery dataset) tested with GAMuT. Adjusted for age at death, education, sex, ancestry, smoking status, post-mortem interval (PMI), 3 or 4 principal components and with or without cell type proportions.

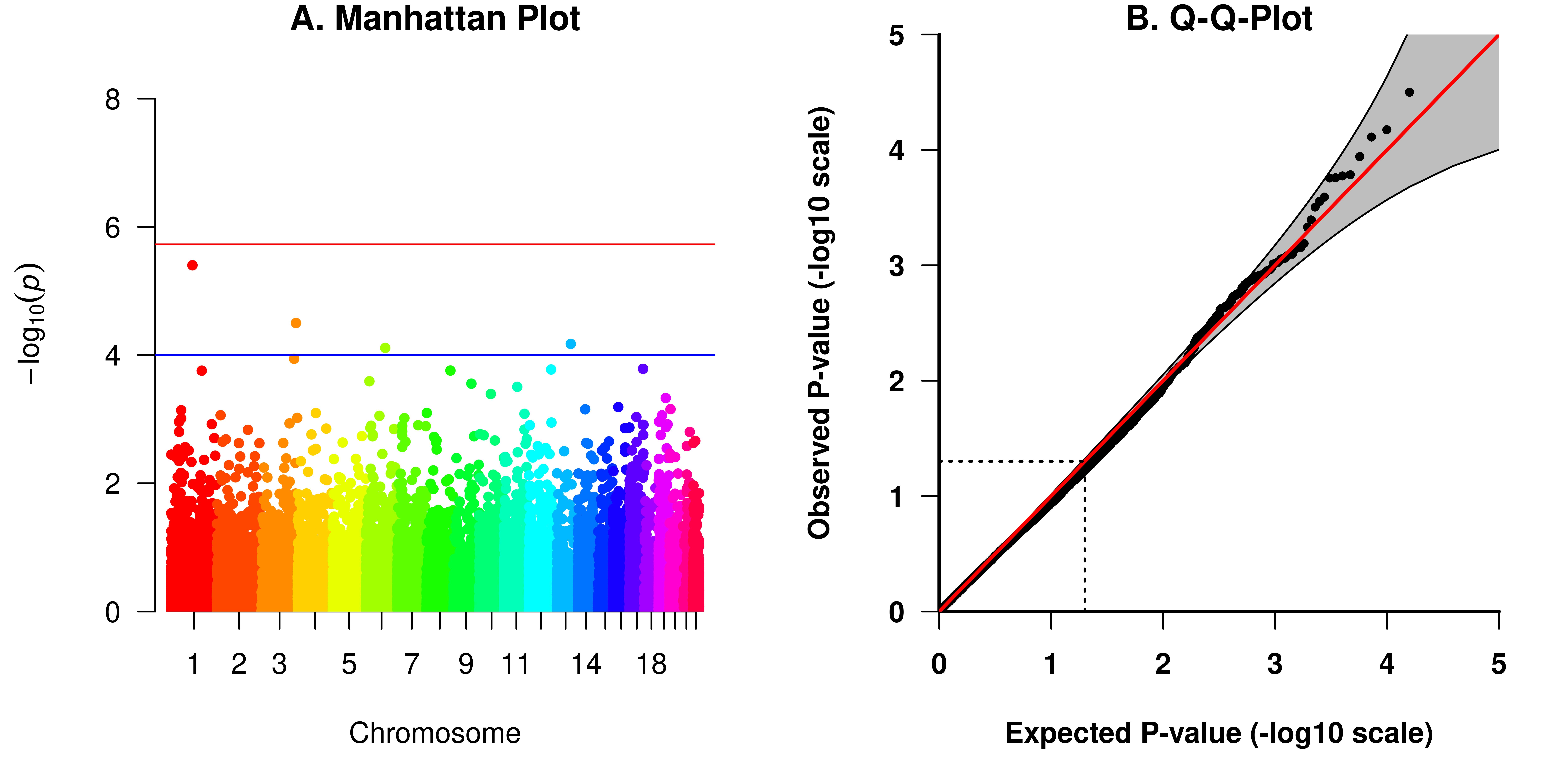

**Figure S6.** Association between DNA methylation and CERAD in ROS/MAP (discovery dataset) tested with GAMuT. Adjusted for age at death, education, sex, ancestry, smoking status, post-mortem interval (PMI), cell type proportions and the first four principal components.

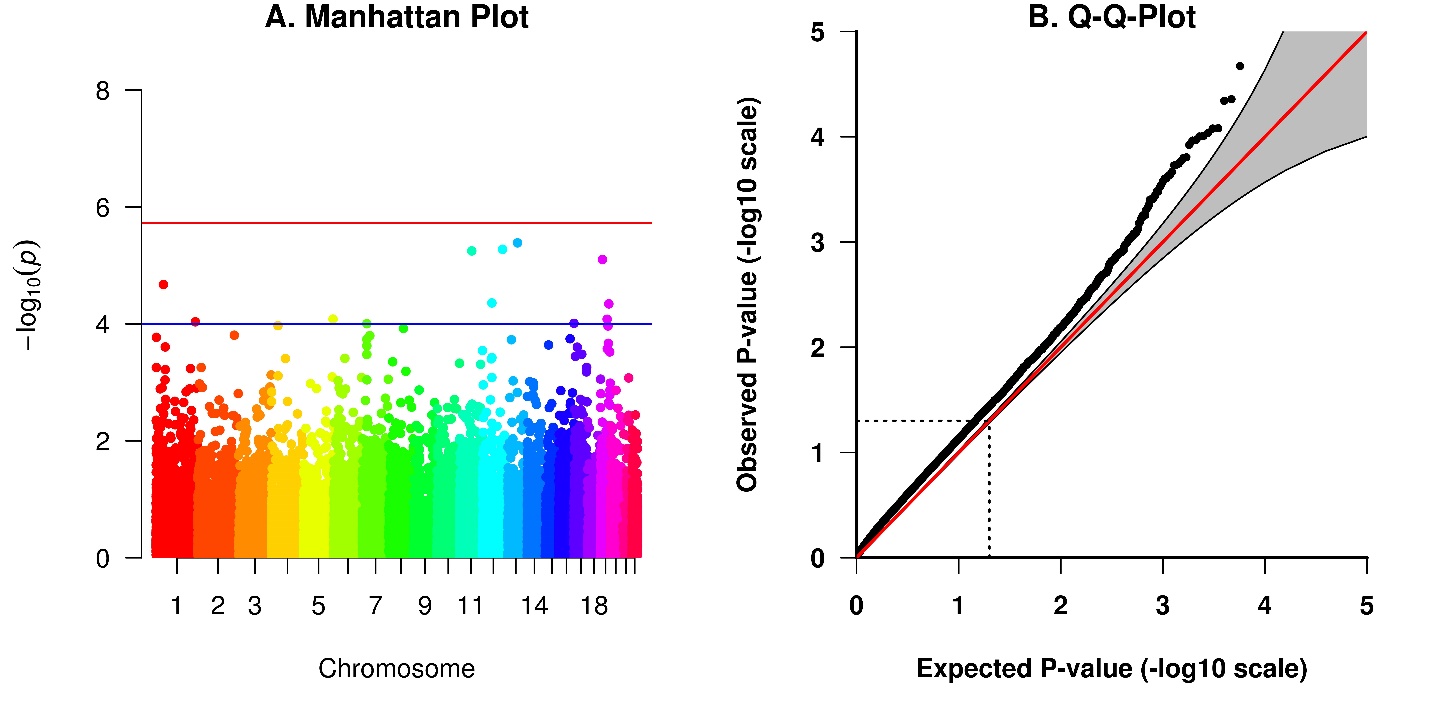

**Figure S7.** Association between DNA methylation and Braak stage in ROS/MAP (discovery dataset) tested with GAMuT. Adjusted for age at death, education, sex, ancestry, smoking status, post-mortem interval (PMI), cell type proportions and the first three principal components.

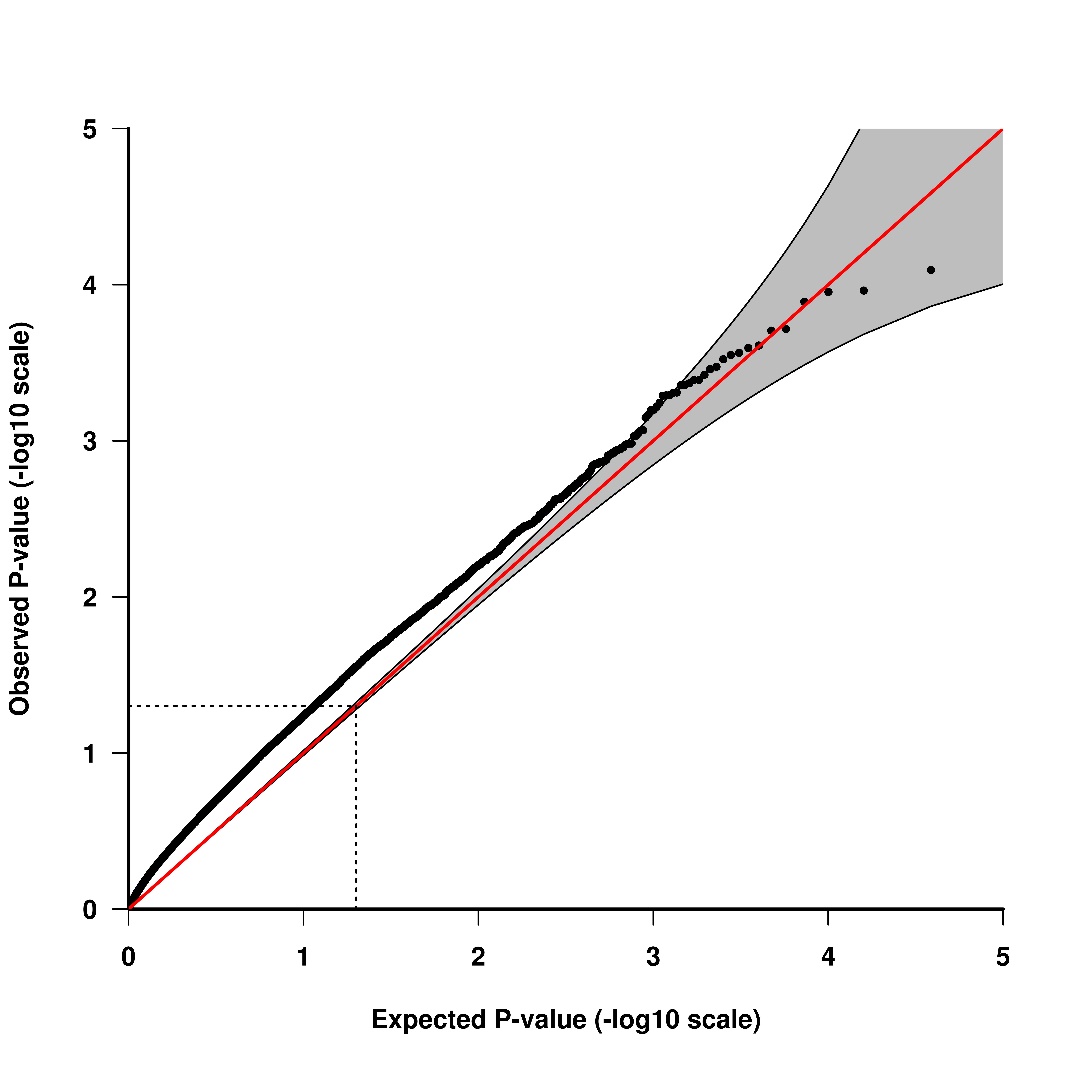

**Figure S8.** Association between DNA methylation and BRAAK stage in the replication dataset (N=66) tested with GAMuT. Adjusted for age at death, sex and cell type composition.

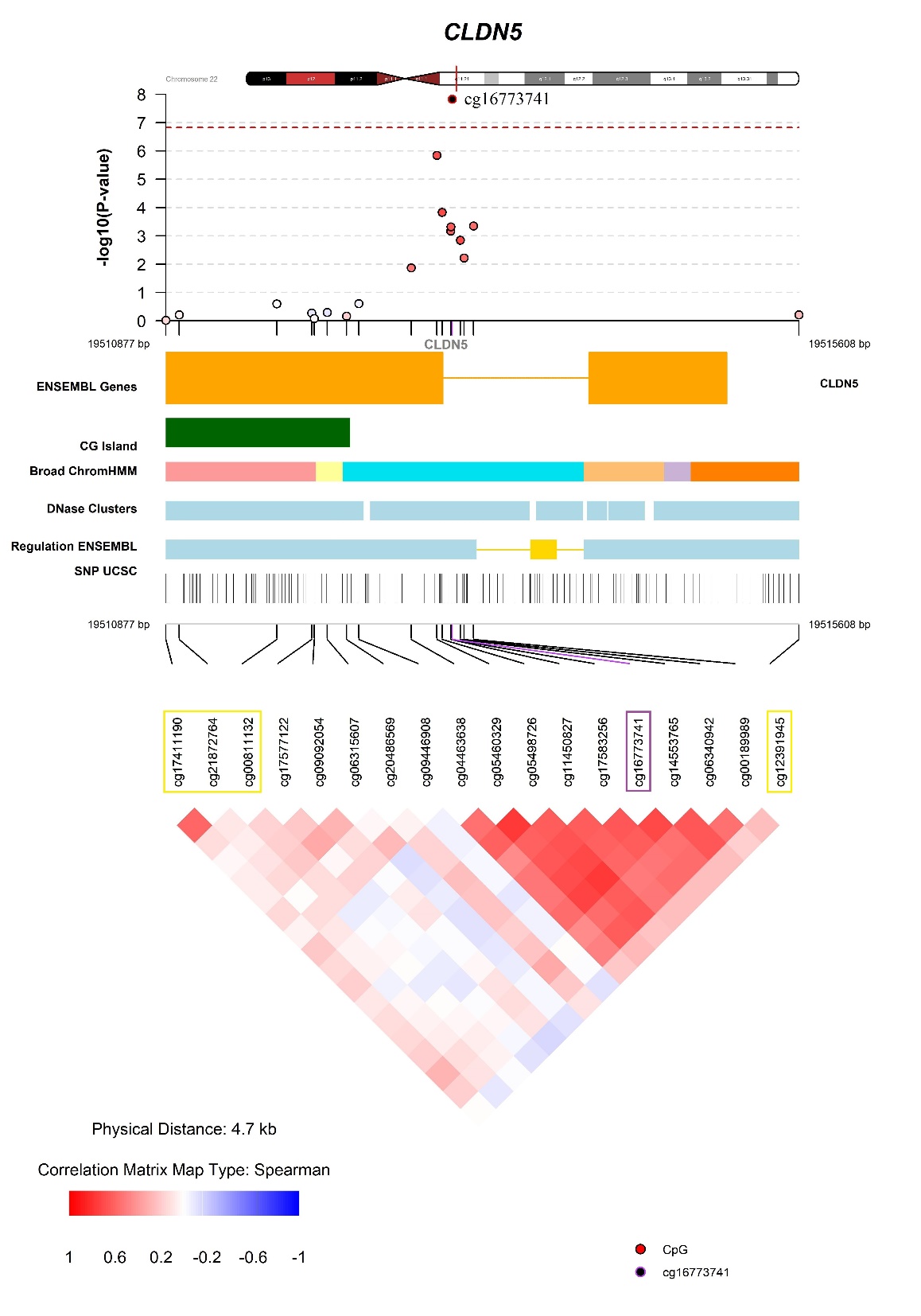

**Figure S9.** Fine mapping of the association between DNA methylation and decline in episodic memory. Results from linear regression analyses on the association between CpG sites and cognitive decline in ROS/MAP (discovery dataset) adjusted for age at death, education, sex, ancestry, smoking status, post-mortem interval (PMI) and the first four principal components. The most significant CpG site (cg16773741) is marked in purple and CpG sites associated with genotypes in the same window are marked in yellow (compare Table S15).

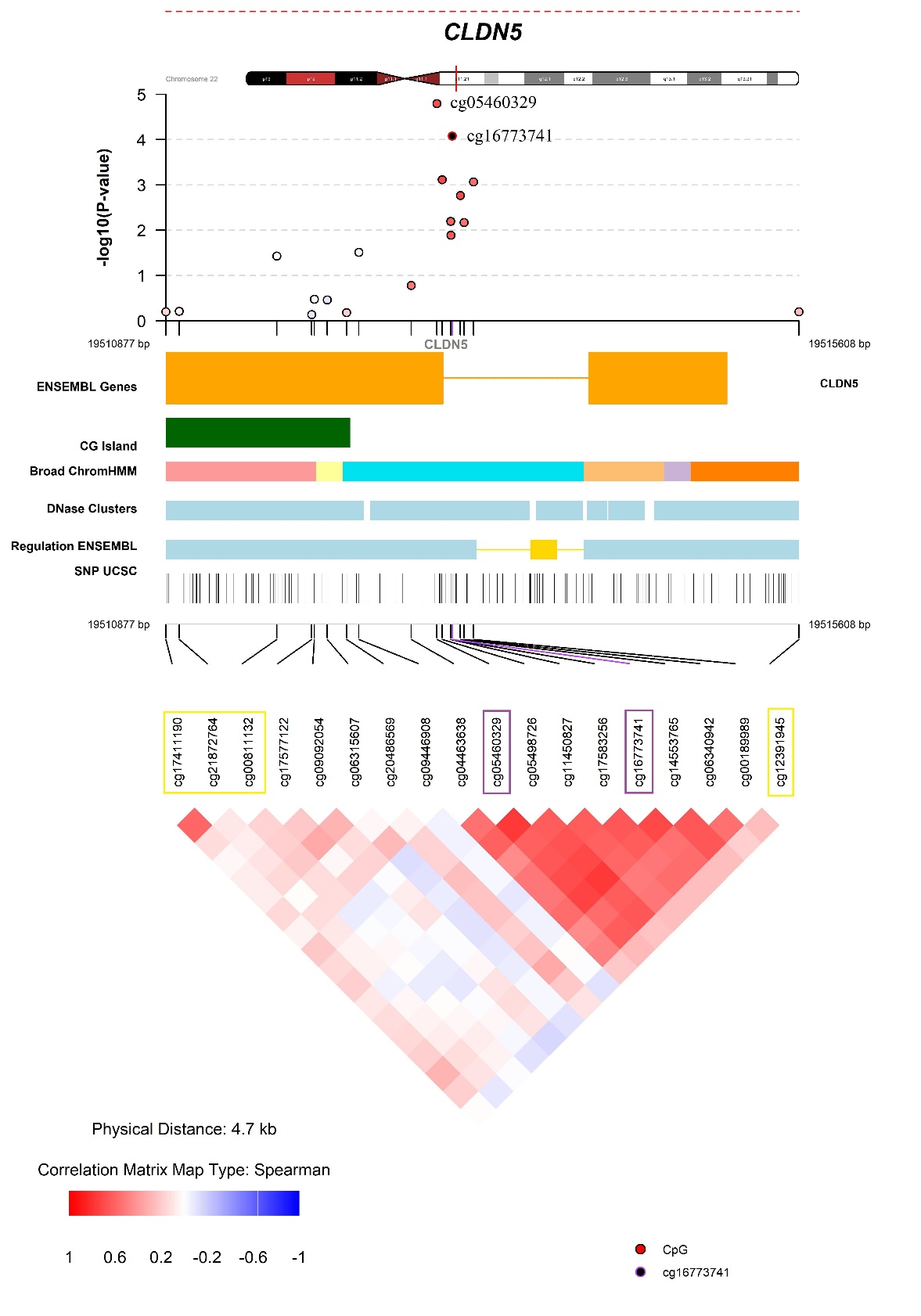

**Figure S10.** Fine mapping of the association between DNA methylation and decline in perceptual speed. Results from linear regression analyses on the association between CpG sites and cognitive decline in ROS/MAP (discovery dataset) adjusted for age at death, education, sex, ancestry, smoking status, post-mortem interval (PMI) and the first four principal components. The most significant CpG site (cg16773741) is marked in purple and CpG sites associated with genotypes in the same window are marked in yellow (compare Table S15).

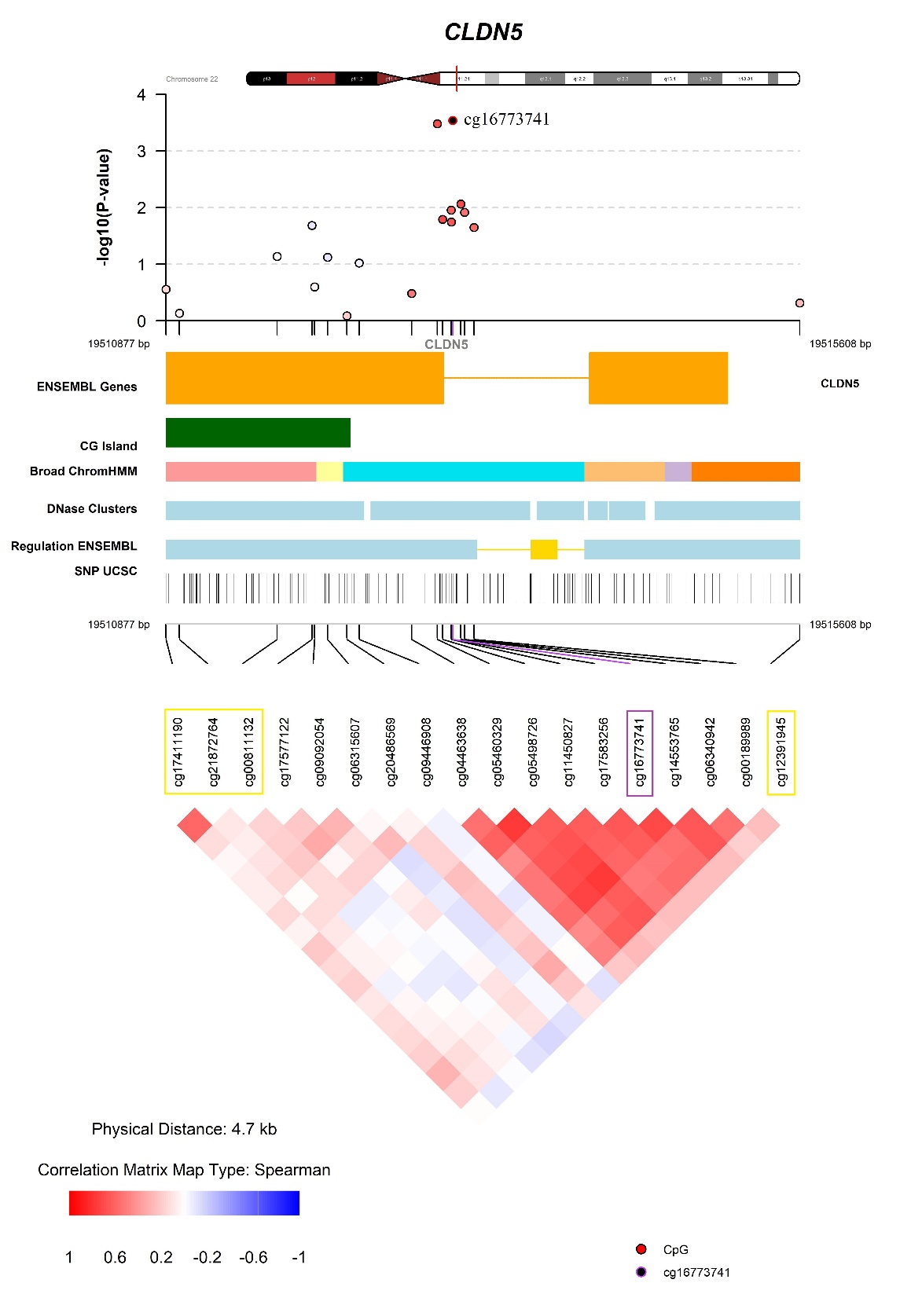

**Figure S11.** Fine mapping of the association between DNA methylation and decline in perceptual orientation. Results from linear regression analyses on the association between CpG sites and cognitive decline in ROS/MAP (discovery dataset) adjusted for age at death, education, sex, ancestry, smoking status, post-mortem interval (PMI) and the first four principal components. The most significant CpG site (cg16773741) is marked in purple and CpG sites associated with genotypes in the same window are marked in yellow (compare Table S15).

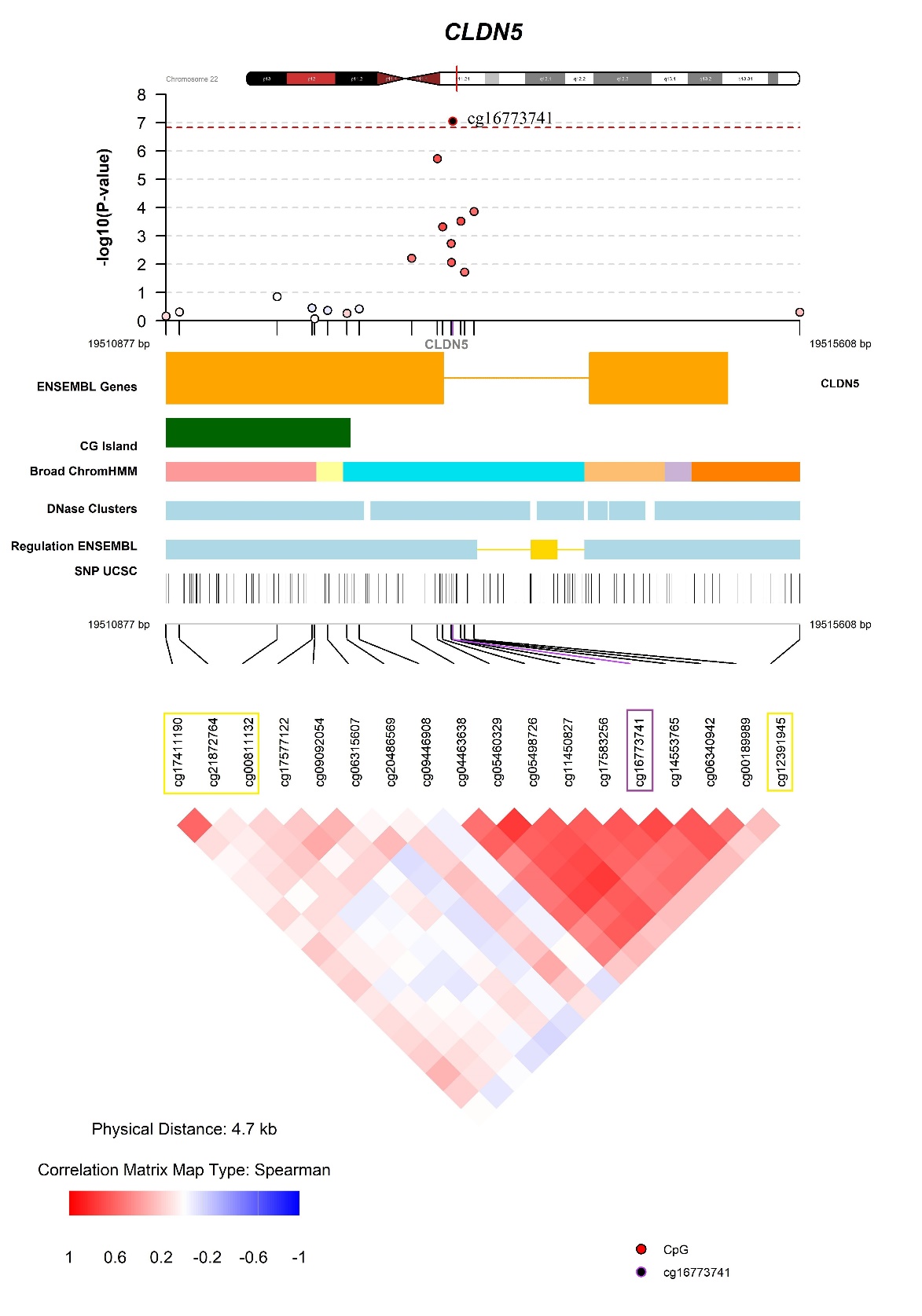

**Figure S12.** Fine mapping of the association between DNA methylation and decline in semantic memory. Results from linear regression analyses on the association between CpG sites and cognitive decline in ROS/MAP (discovery dataset) adjusted for age at death, education, sex, ancestry, smoking status, post-mortem interval (PMI) and the first four principal components. The most significant CpG site (cg16773741) is marked in purple and CpG sites associated with genotypes in the same window are marked in yellow (compare Table S15).
